# Acetylation of lysine 82 initiates TDP-43 nuclear loss of function by disrupting its nuclear import

**DOI:** 10.1101/2024.09.04.611121

**Authors:** Sitao Zhang, Sonia Vazquez-Sanchez, Shan Lu, Michael W. Baughn, Jaisen Lim, Spencer Oung, Lilian Gao, Jolene K. Diedrich, John R Yates, Haiyang Yu, John Ravits, Don W. Cleveland

## Abstract

The hallmark of a spectrum of age-dependent neurodegenerative diseases, including Amyotrophic Lateral Sclerosis (ALS), is a TDP-43 proteinopathy that includes nuclear loss of function and cytoplasmic aggregation. Here, reduced proteasome activity, as naturally occurs during aging, is shown to inhibit nuclear import of TDP-43. Quantitative mass spectrometry is used to determine that TDP-43 is the protein whose nuclear localization is most perturbed upon reduction in proteasome activity, culminating in elevated cytoplasmic TDP-43. Interaction of importin-α1 with the bipartite <u>c</u>lassical <u>n</u>uclear localization <u>s</u>equence (cNLS) of TDP-43 is shown to be disrupted by partial proteasome inhibition but maintained by replacement with a PY-NLS that is recognized by importin-β2. Mechanistically, this nuclear depletion of TDP-43 is shown to be driven by ubiquitination or acetylation of lysines 79, 82, and 84 within the cNLS when proteasome activity is reduced in human neurons. Specifically, acetylation at lysine 82 is sufficient to abolish TDP-43 binding to importin-α1 and subsequent nuclear import of TDP-43. Moreover, using acetylation-specific TDP-43 antibodies, we detected acetylation of lysine 82 in the motor cortex of sporadic ALS patients but not control subjects. Our findings demonstrate that post-translational acetylation at lysine 82 of TDP-43 drives disruption of its importin-α1-mediated nuclear import and is sufficient to initiate TDP-43 nuclear loss of function and cytoplasmic accumulation, evidence supporting acetylation as a plausible initiator of TDP-43 proteinopathies.

## Introduction

Loss of nuclear TDP-43 (<u>T</u>AR <u>D</u>NA/RNA-binding <u>p</u>rotein <u>43</u> kDa) and its aggregation in the cytoplasm, known as TDP-43 proteinopathy, is a neuropathological feature found in 97% of ALS cases and in many other age-dependent neurodegenerative diseases, including Frontotemporal dementia (FTD)^1^, Alzheimer’s disease (AD)^2–6^, and <u>L</u>imbic-predominant <u>a</u>ge-related <u>T</u>DP-43 <u>e</u>ncephalopathy (LATE)^7^. LATE TDP-43 proteinopathy appears in brains of 20-50% of people older than 80 years and is estimated to be 100-fold more frequent than FTD^7^. In these neurodegenerative diseases, neurons lose nuclear TDP-43, in turn accumulating it (or fragments^8^ of it) in hyperphosphorylated^8^, ubiquitinated^8^, and acetylated^9^ cytoplasmic aggregates. Missense mutations in TDP-43 are associated with familial FTD and ALS^10–13^ indicating a causative link.

The normal functions of the predominantly nuclear TDP-43 include RNA transcription, splicing, and transport^14^. TDP-43 is functional in both the nucleus and the cytoplasm and, therefore, its trafficking across the nuclear membrane is essential for its proper function^14^. TDP-43 binds to RNA with high specificity via its two RNA recognition motifs (RRMs)^14^. The nuclear export of TDP-43 is thought to be achieved by passive diffusion, as its predicted nuclear export signals^15^ have been shown to be neither necessary nor sufficient for its nuclear export^16–18^. Nuclear import of TDP-43 is facilitated by its bipartite classical nuclear localization sequence (cNLS) that is recognized by the importin-α/importin-β1 (also known as KPNA/KPNB1) import pathway^19^. Molecular mechanisms that modulate TDP-43 localization to the nucleus and cytoplasm and their contribution to TDP-43 proteinopathy are not known.

Here, we identify a mechanism that is induced by age-related decrease in proteasome activity and leads to the nuclear depletion and cytoplasmic accumulation of TDP-43 in cultured human neurons. Furthermore, we determine that this molecular pathway is used in the motor cortex of individuals with sporadic ALS. Specifically, we show that partial reduction in proteasome activity inhibits TDP-43 nuclear localization through ubiquitination and acetylation within its cNLS, the latter modification governed by competing activities of acetyltransferases and deacetylases. Acetylation-mimicking modification of lysine K82 (and to a lesser extent K79 and K84) that lies within the TDP-43 cNLS is shown to almost completely block TDP-43 nuclear import in cultured human cortical neurons. Moreover, acetylation of lysine 82 within the TDP-43 cNLS is found in postmortem samples from sporadic ALS, consistent with posttranslational modifications of the TDP-43 NLS inducing TDP-43 cytoplasmic accumulation (and corresponding nuclear depletion) as a plausible initiator of TDP-43 proteinopathy.

## Results

### TDP-43 mislocalization is triggered by reduced proteasome activity

Proteasome activity has been claimed to decline during aging of metazoans^20–22^. Using lysates prepared from fresh frozen cortex of mice of different ages, we confirmed a ∼50% decline in chymotrypsin-like proteasome activity by one year of age ***(Fig. S1A).*** Similarly, we found a 20-60% decline in the chymotrypsin-like proteasome activity in cell lysates from fresh frozen motor cortices in all sporadic ALS samples tested ***(Fig. S1A).*** To mimic this age-dependent reduction in proteasome activity, we determined the levels needed of three well characterized proteasome inhibitors bortezomib [BTZ, a reversible inhibitor of the chymotrypsin-like activity of the proteasome and used as an FDA approved anticancer drug for refractory multiple myeloma^23^]), MG132, or marizomib [MRZ], to achieve ∼50% proteasome inhibition in induced pluripotent stem cell (iPSC)-derived human cortical neurons (2 nM, 100 nM, and 10 nM, respectively) (***Fig. 1A, Fig. S1B***). Neuronal viability was unaffected after 1 week (***Fig. S1C***) of reduced proteasome activity.

**Figure 1:**
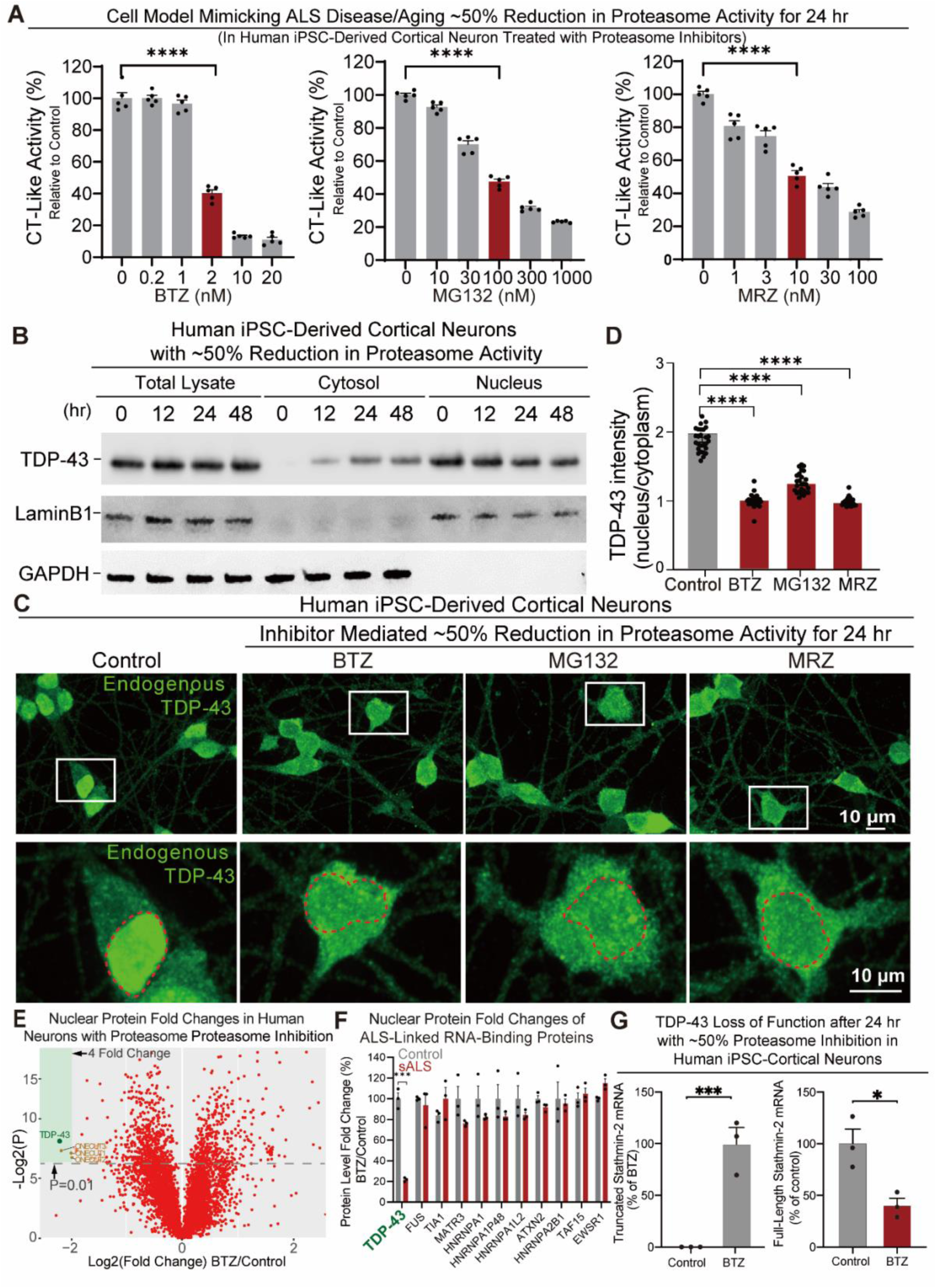
Partial reduction in proteasome activity results in TDP-43 mislocalization and loss of function. **(A)** Chymotrypsin-like [(CT)-like] proteasome activity in cell lysates from human iPSC-derived cortical neurons after exposure for 24 hr to varying doses of proteasome inhibitors BTZ, MG132, and MRZ. **(B)** TDP-43 protein levels determined by immunoblotting total and fractionated (nucleus and cytosol) lysates of human iPSC-derived cortical neurons with or without BTZ (2 nM) exposure for 0, 12, 24 and 48 hr. Corresponding immunoblots for Lamin B1 and GAPDH provide loading controls for nuclear fraction and cytosolic fractions, respectively. (**C)** Immunofluorescent examples of endogenous TDP-43 (Green) in human iPSC-derived cortical neurons after 24 hr of treatment with proteasome inhibitors (BTZ: 20 nM, MG132: 100 nM, MRZ: 100 nM). **(D)** Quantification of the nucleocytoplasmic ratio of TDP-43 in images from Fig. 1D. **(E)** Volcano plot of statistical significance versus the log2 fold change for the 5202 proteins in the nuclear fraction of human iPSC-derived cortical neurons after 24 hr (or not) of exposure to 20 nM BTZ. Unadjusted P values were calculated using a one-sample two-sided Student’s t-test. **(F)** Quantification of nuclear levels of ALS-linked RNA-binding proteins (in Fig. 1E) in human iPSC-derived cortical neurons with or without exposure to BTZ for 24 hr. **(G)** RT-PCR analysis of full-length and truncated stathmin-2 mRNA levels in human iPSC-derived cortical neurons with or without exposure to 20 nM BTZ for 24 hr.

In these *in vitro* neuronal models, we assessed TDP-43 localization after a reduction in proteasome activity similar to that observed upon aging and ALS. At 0, 12, 24, or 48 hr after 50% inhibition of proteasome activity in fractionated cell lysates the total TDP-43 protein level was unchanged, yet as early as 12 hr after reduction in proteasome activity TDP-43 was cytoplasmically mislocalized (as shown by immunoblotting of fractionated neuronal cell lysates- ***Fig. 1B; Fig. S1D***). Indeed, analysis with indirect immunofluorescence confirmed increased cytosolic TDP-43 contemporaneous with decreased nuclear TDP-43 following proteasome inhibition by 50% for 24 hr with BTZ, as well as with MG132 and MRZ (***Fig. 1B-E; Fig. S1E,F***).

TDP-43 mislocalization was followed by real-time live-cell imaging of iPSC-derived cortical neurons transduced to express fluorescently tagged TDP-43 (TDP-43-Clover) with (by addition of 2 nM BTZ) or without proteasome inhibition. While TDP-43-Clover was almost exclusively nuclear prior to BTZ addition, within 12 hr TDP-43-Clover mislocalization was evident, with approximately half of TDP-43 mislocalized within 24 hr (***Fig. S1G***). Immunoblotting of cell fractions from human iPSC-derived cortical neurons, either untreated or treated with BTZ (2 nM) for 24 hr, confirmed that both endogenous TDP-43 and TDP-43-Clover became more cytoplasmic with similar kinetics (***Fig. S1H***). Taken together, TDP-43 accumulates in the cytoplasm of cortical human neurons following partial reduction in proteasomal activity.

### TDP-43 is the protein whose nuclear localization is most sensitive to reduced proteasome activity

To determine if TDP-43 mislocalization in neurons upon proteasome inhibition was specific to TDP-43 or a more general effect on nuclear import, we used quantitative Tandem Mass Tag (TMT) mass spectrometry to determine the nuclear proteome of the iPSC-derived cortical neurons with or without BTZ-induced proteasome inhibition. Remarkably, TDP-43 was the protein whose nuclear localization was the most sensitive to a reduction in proteasome activity, with its nuclear content reduced by more than 4 fold within 24 hr (***Fig. 1E,F***). Partial nuclear clearance was selective for TDP-43, as nuclear content of other ALS-linked RNA-binding proteins was not (or barely) affected **(*Fig. 1F*)**.

To test whether this mislocalization affected TDP-43 nuclear function, we assessed the usage of cryptic splicing/polyadenylation sites within TDP-43-regulated stathmin-2 pre-mRNAs. Indeed, the TDP-43 mislocalization was sufficient to induce cryptic splicing of stathmin-2 as it 1) generated non-productive, truncated stathmin-2 mRNA and 2) reduced the level of full-length stathmin-2 mRNA (***Fig. 1G***), as demonstrated following reduced TDP-43 function^24,25^. This TDP-43 loss of function also corresponded with doubling of the level of TDP-43 encoding RNAs (***Fig. S1I***), in line with dysfunction of the known TDP-43 autoregulation pathway when nuclear TDP-43 is reduced^26,27^.

### Reduced proteasome activity disrupts importin-α1 recognition of TDP-43

To test if 24 hr of decreased proteasome activity inhibited TDP-43 interaction with nuclear import receptors, we immunoprecipitated TDP-43 and addressed its binding to importin proteins (importin-α1 and importin-β2) (see scheme in ***Fig. S2A***). While a proportion of importin-α1, but not importin-β2, was initially bound to TDP-43 in iPSC-derived neurons, proteasome inhibition completely disrupted interaction with importin-α1 (***Fig. 2A***), but did not affect binding of the ALS-linked multifunctional DNA/RNA-binding protein FUS (Fused in Sarcoma) to its nuclear import partner importin-β2 that recognizes the non-classical PY-NLS of FUS^28^ (***Fig. 2B***), nor FUS’s primarily nuclear localization (***Fig. 2C***). Immunoblot analysis of TDP-43 and FUS protein levels in total cell lysate, nucleus, and cytosol samples from human iPSC-derived cortical neurons, either untreated or treated with BTZ (2 nM) for 24 hr, confirmed that TDP-43, but not FUS, exhibited increased cytoplasmic mislocalization when proteasome activity was reduced (***Fig. S2B;* *Fig. 1F,E***).

**Figure 2:**
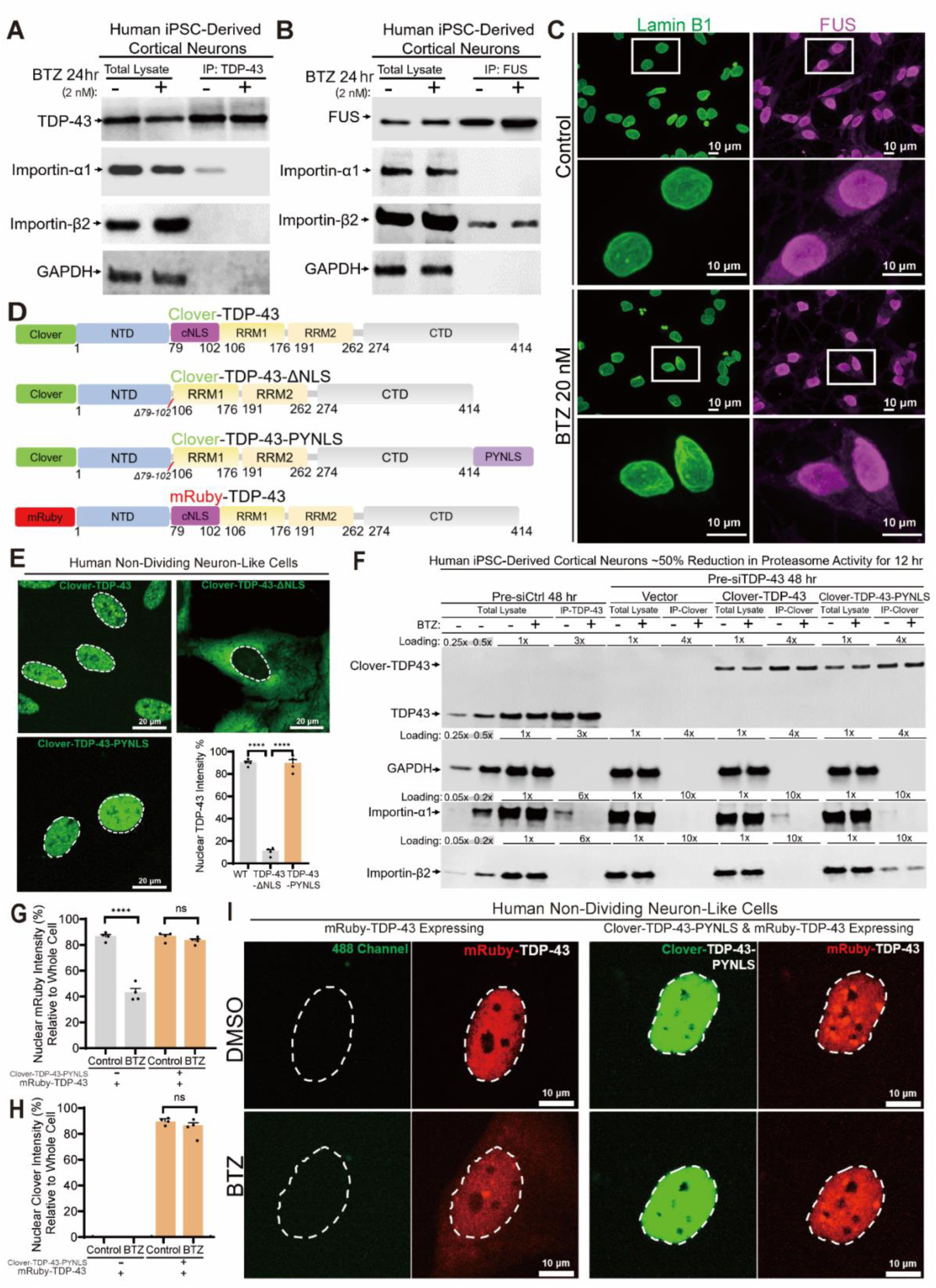
The classical NLS (cNLS) of TDP-43 mediates TDP-43 mislocalization upon partial proteasome inhibition that is rescued by swapping the TDP-43 cNLS with the FUS-PY-NLS. **(A)** Co-immunoprecipitation (co-IP) was conducted with a primary antibody for TDP-43 in iPSC-derived cortical neurons after 24 hr of proteasome inhibition (2 nM BTZ) and control. Immunoblot for TDP-43, importin-α1, importin-α5, and importin-β2. GAPDH was used as a loading control. **(B)** Co-immunoprecipitation (co-IP) was conducted with a primary antibody for FUS in iPSC-derived cortical neurons after 24h of proteasome inhibition (2 nM BTZ) and control. Immunoblot to determine levels of TDP-43, importin-α1, importin-α5, and importin-β2. GAPDH was used as a loading control. **(C)** Representative images using confocal microscopy of iPSC-derived cortical neurons after 24 hr of proteasome inhibition (20 nM BTZ) and control immunostained with Lamin B1 (green) and FUS (magenta). **(D)** Schematic of TDP-43 WT, TDP-43-ΔNLS, and TDP-43-PYNLS in which the cNLS of TDP-43 was replaced by the PY-NLS of FUS. All proteins were tagged at the N terminus with Clover (a bright GFP variant) or mRuby, respectively. **(E)** Representative live-cell images of Clover-TDP-43, Clover-TDP-43-ΔNLS, and Clover-TDP-43-PY-NLS localization and quantification in non-dividing, human neuron-like SH-SY5Y cells. **(F)** Co-immunoprecipitations (co-IP) were conducted with TDP-43 antibody-coated magnetic beads and GFP nanobody coated magnetic beads in cell lysates of correspondingly tagged TDP-43 in iPSC-derived cortical neurons. Neurons were pre-treated with either control or human TDP-43 siRNA for 48 hr, followed with transduction for 72 hr with lentiviruses encoding Clover-TDP-43-WT or Clover-TDP-43-PY-NLS. Each cell group was then treated with (or not) 2 nM BTZ for an additional 12 hr. Immunoblot shows levels of TDP-43, importin-α1, and importin-β2. GAPDH was used as a loading control. **(G)** Quantification of nuclear versus whole cell mRuby fluorescence intensity in human neuron-like SH-SY5Y cells stably expressing mRuby-TDP-43-WT alone or co-expressing with Clover-TDP-43-PY-NLS with or without partial proteasome inhibition induced with BTZ (2 nM) for 0-6 hr. **(H)** Quantification of nuclear versus total cell fluorescence intensity for Clover-TDP-43-PYNLS in human SH-SY5Y cells stably expressing mRuby-TDP-43-WT alone or co-expressing with Clover-TDP-43-PYNLS with or without partial BTZ (2 nM)-induced proteasome inhibition for 0-6 hr. **(I)** Representative live-cell images of mRuby-TDP-43 (alone) and both mRuby-TDP-43 & Clover-TDP-43-PY-NLS in non-dividing SH-SY5Y cells with or without BTZ (2 nM)-induced proteasome inhibition for 0-6 hr.

Next, we used live-cell imaging to follow Clover-tagged TDP-43 localization upon 50% proteasome inhibition (***Fig. 2D,E***). Replacing the TDP-43 cNLS with the FUS-derived PY-NLS in Clover-tagged TDP-43 was sufficient to generate TDP-43 which was bound by importin-β2 instead of importin-α1 (***Fig. 2F***) and to abrogate TDP-43 mislocalization when proteasome activity was reduced (***Fig. 2E***). Using dual color tags of TDP-43 and a proteasome inhibitor (2 nM BTZ) for 0-6 hr (***Fig. S2C***), we found that Clover-TDP-43-PY-NLS was not only resistant to mislocalization, but also prevented cNLS-containing, mRuby-tagged TDP-43 from mislocalization (***Fig. 2G-I***), probably through N-terminal dimerization or oligomerization of PY-NLS- and cNLS-containing TDP-43 molecules during importin-β2 mediated import or inside the nucleus^29^.

Immunoblotting of neuronal cell lysates confirmed that Clover-TDP-43-PY-NLS subcellular localization was resistant to a 50% decrease in proteasome activity from BTZ exposure for 24 hr (***Fig. S2D***). Furthermore, substitution of the TDP-43 cNLS with the FUS PY-NLS prevented TDP-43 loss of nuclear function in human neurons, as it sustained stathmin-2 protein levels observed after reduction in proteasome activity (***Fig. S2E***). Collectively, these data indicate that the cNLS of TDP-43 is pivotal to its mislocalization upon a mild decrease in proteasomal activity.

### Mislocalization of TDP-43 from modification of the TDP-43 cNLS

Following editing of both endogenous alleles to produce full length TDP-43 with a C-terminal Clover tag, TDP-43 was affinity purified (***Fig. S3A***) from non-dividing human neuron-like cells which had been treated for 24 hr (or not) with BTZ to partially inhibit proteasome activity. Post-translational modifications were then identified with mass spectrometry after proteasome inhibition (***Fig. 3A***). With nearly complete (98.3%) peptide coverage, including the complete bipartite cNLS (***Fig. S3B***), acetylation and/or ubiquitination of lysines 79, 82, and 84 were identified within the cNLS, as well as phosphorylation of serines 91 and 92 (***Fig. 3B***).

**Figure 3:**
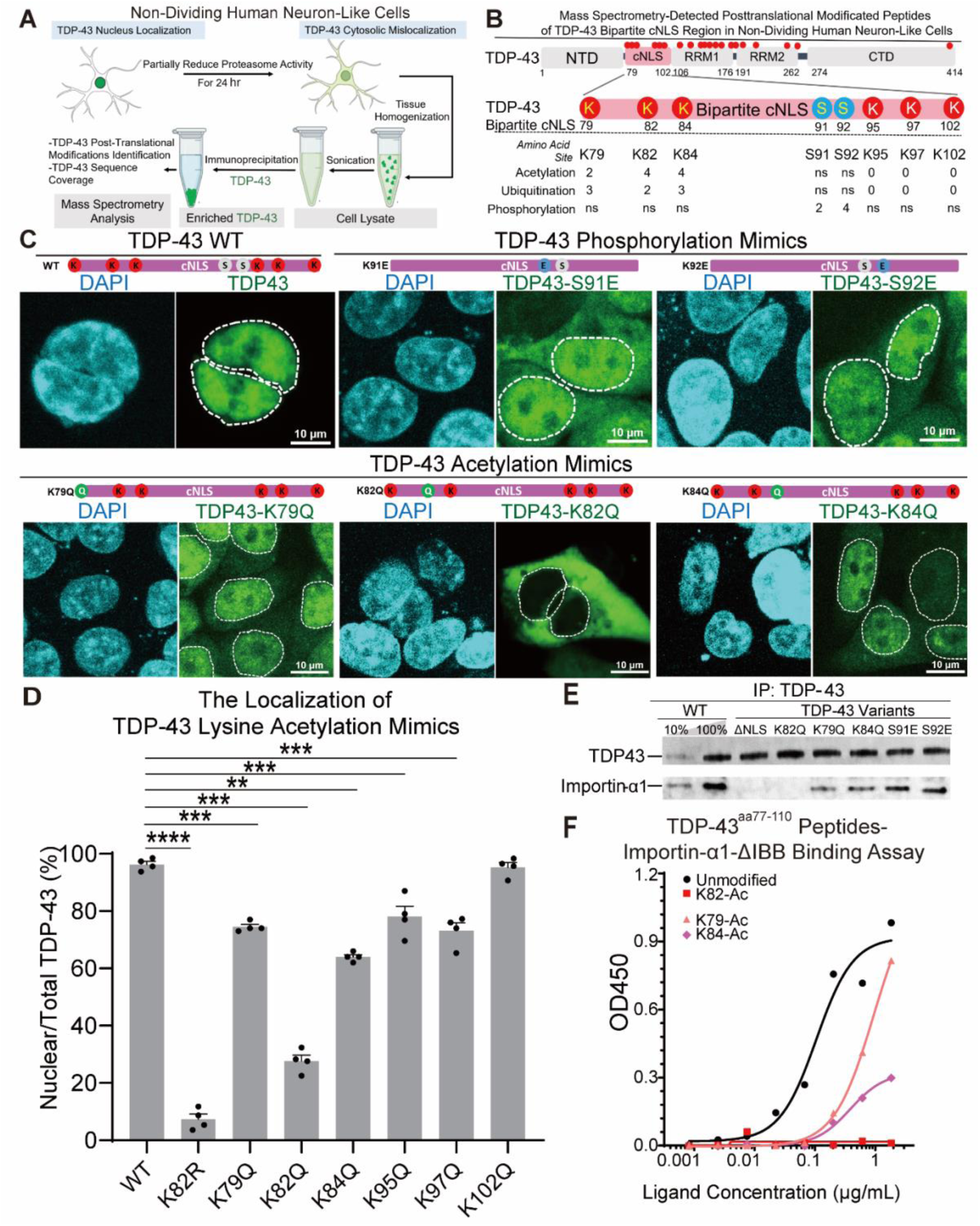
Post-translational acetylation, ubiquitination, and phosphorylation within the TDP-43 cNLS are induced by partial proteasome inhibition and mimicking them disrupts the TDP-43-importin-α1 interaction driving TDP-43 mislocalization. **(A)** Scheme for detection of post-translational modification of TDP-43-Clover purified from non-dividing human neuron-like SH-SY5Y cells with homozygous knock-in of TDP-43-Clover at both endogenous TDP-43 alleles. SH-SY5Y cells with or without proteasome inhibitor BTZ (2 nM) for 24 hr were harvested and lysed. The TDP-43-Clover proteins were collected using GFP nanobodies conjugated to magnetic beads. After elution, mass spectrometry was used to identify post-translational modifications. **(B)** Scheme for detecting post-translational modifications within the cNLS of TDP-43 identified from purified TDP-43 samples as described in (A). **(C)** Representative live-cell fluorescence images of C-terminally Clover tagged TDP-43-WT, TDP-43 phosphorylation mimics (TDP-43-S91E and TDP-43-S92E), and TDP-43 acetylation mimics (TDP-43-K79Q, TDP-43-K82Q, and TDP-43-K84Q) expressed in non-dividing human neuron-like SH-SY5Y cells after pre-treatment with either control or human TDP-43 siRNA for 48 hr, followed with lentivirus-mediated expression of different TDP-43 variants with C-terminal Clover tag for 72 hr. **(D)** Quantification of nuclear TDP-43-Clover variant levels in (C). **(E)** Co-immunoprecipitation (co-IP) with GFP nanobody-magnetic beads of cell lysates from SHSY-5Y cells pre-treated with human TDP-43 siRNA for 48 hr, followed by lentivirus-mediated expression for 72 hr of C-terminal Clover tagged TDP-43 modification mimics in (C). Levels of TDP-43 and importin-α1 are determined by immunoblot. **(F)** TDP-43 peptide-importin-α1 binding assay conducted across a range of concentrations with importin-α1-ΔIBB and 33 amino acid peptides (TDP-43^aa77–110^) synthesized with or without acetylation at K79, K82, or K84, and phosphorylation at S92. Random peptides were used as a negative control; PBS was used as a blank control.

The consequence of each of these modifications within the TDP-43 cNLS on nuclear import was determined using lentiviral transduction of iPSC-derived cortical neurons to express wild-type TDP-43 or variants of it containing acetylation-(lysine to glutamine) or phosphorylation (serine to glutamic acid) mimicking substitutions at the sites identified to be post-translationally modified. Endogenous TDP-43 was removed by a 48 hr pre-treatment with human TDP-43 siRNAs (***Fig. 2F, S2E***), followed by 72 hr of lentivirus-mediated expression of the TDP-43 variants (see schematic in ***Fig. S3C***). While phosphorylation (S91E and S92E) and acetylation (K79Q and K84Q) mimics of TDP-43 modestly affected TDP-43 nuclear content and partially disrupted importin-α1 binding, incorporation of the acetylation-mimicking variant TDP-43-K82Q eliminated its nuclear import and binding to importin-α1 (***Fig. 3C-E***), consistent with a recent report^30^.

To further test whether actual posttranslational modification at or near lysine 82 affected TDP-43-importin-α1 interaction, we determined the binding capacities to importin-α1 of TDP-43^aa77–110^ peptides with or without acetylation at K79, K82, or K84, and phosphorylation at S92. Peptides acetylated at K82 did not bind to importin-α1, while peptides acetylated at K79 or K84, as well as those phosphorylated at S92, exhibited reduced binding to importin-α1 (***Fig. 3F***). Taken together, reduced proteasome activity induces acetylation, and to a lesser extent phosphorylation, of residues – particularly K82 – within the TDP-43 cNLS. This disrupts binding to importin-α1, causing failure of TDP-43 nuclear import.

### Lysine 82 in the cNLS is a gatekeeper of TDP-43 nuclear import

Point mutations in the TDP-43 cNLS were generated and expressed in human neuron-like cells with siRNA-mediated TDP-43 depletion to pinpoint the determinants of TDP-43 central to its interaction with importin-α1. Replacing all 6 lysines in the TDP-43 cNLS with arginines (the most conservative point mutation of lysine and one that retains its positive charge) to produce TDP-43-6KR resulted in complete cytosolic localization (***Fig. 4A,B***), while replacing all other lysines outside the NLS (TDP-43-14KR) did not affect TDP-43 nuclear localization (***Fig. 4A,B***). Evaluation of individual changes at each of the lysines in the TDP-43 cNLS (***Fig. 4A***) revealed that TDP-43-K82R (but not the other variants) induced TDP-43 mislocalization (***Fig. 4D,E***) even when accumulated at only 25% the level of endogenous TDP-43 (***Fig. S4A***). Correspondingly, the TDP-43/importin-α1 interaction was abrogated for TDP-43-6KR and TDP-43-K82R (***Fig. 4C***), findings that support TDP-43 lysine 82 to be critical for the TDP-43-importin-α1 interaction essential for nuclear TDP-43 localization.

**Figure 4:**
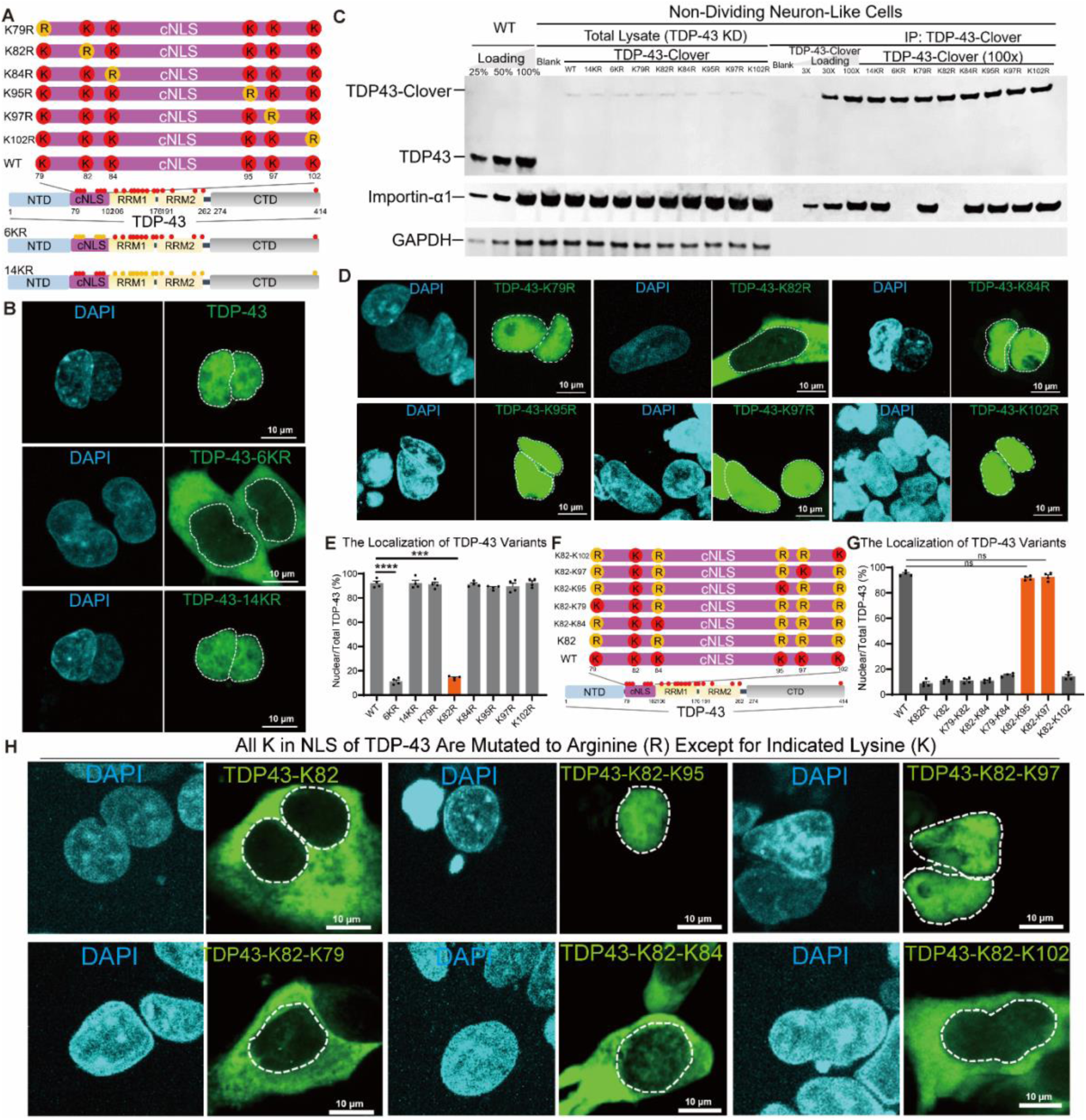
Lysine 82, situated within the cNLS of TDP-43, plays a central role in regulating the nuclear localization of TDP-43. **(A)** Schematic of TDP-43 NLS mutations. A Clover tag was placed at C-terminus of each variant. **(B)** Representative live-cell images of TDP-43-Clover, TDP-43-6KR-Clover, and TDP-43-14KR-Clover in non-dividing human neuron-like SH-SY5Y cells pre-treated with either control or human TDP-43 siRNA for 48 hr, followed with lentivirus-mediated expression for 72 hr of the different Clover tagged TDP-43 variants. **(C)** Co-immunoprecipitation (co-IP) with GFP nanobody-magnetic beads of lysates of non-dividing SHSY-5Y cells after pre-treatment with human TDP-43 siRNA for 48 hr, followed with lentivirus-mediated expression for 72 hr of Clover-TDP-43-WT or Clover-TDP-43-PY-NLS. TDP-43, importin-α1, and importin-β2 levels were determined by immunoblot. GAPDH was used as a loading control. **(D)** Representative live-cell images of Clover-tagged TDP-43 variants harboring cNLS point mutations in non-dividing SH-SY5Y cells pre-treated with either control or human TDP-43 siRNA for 48 hr, followed with lentivirus-mediated expression for 72 hr of different TDP-43 variants with C-terminal Clover tags. **(E)** Quantification of fluorescence intensity of nuclear TDP-43 in (B,D). **(F)** Schematic of TDP-43 wild-type and lysine to arginine variants in the TDP-43 cNLS. Initially, variant called K82 was produced by converting all Ks except K82 to R. Then, each of the other Ks was reverted from R to K producing a set of variants with only 2 lysines intact to produce: K79-K82, K82-K84, K79-K84, K82-K95, K82-K97, and K82-K102. All proteins were tagged with Clover at the C-terminus. **(G)** Quantification of nuclear TDP-43 fluorescence of each TDP-43 variant in non-dividing human SHSY-5Y cells and **(H)** corresponding, representative live-cell images.

However, although necessary for TDP-43 nuclear import, K82 was not sufficient, as a TDP-43 variant retaining K82 but in which the other 5 lysines within the TDP-43 cNLS were converted to arginine (***Fig. 4F***) was not efficiently imported (***Fig. 4G&H***). Analysis of variants containing K82 and one additional lysine (e.g., K79-K82, K82-K84, K82-K95, K82-K97, and K82-K102) identified that either intact K82-K95 and K82-K97, but not K79-K82, K82-84 or K82-K102 maintained full interaction with importin-α1 at a level similar to wild-type TDP-43 (***Fig. S4B***) and maintained TDP-43 nuclear localization (***Fig. 4G&H***). Most notably, K82 and either K95 or K97 were necessary for TDP-43/importin-α1 interaction and subsequent TDP-43 nuclear import, but acetylation at K82 alone was sufficient to drive complete TDP-43 nuclear clearance.

### Acetylation of TDP-43 on lysine 82 in postmortem motor cortices of sALS patients

To test whether TDP-43 lysine acetylation increases in nervous system samples from sALS, we immunopurified TDP-43 from lysates of fresh frozen human motor cortices (***Fig. 3F***). Since lysine 82 is critical for TDP-43 nuclear localization (***Fig. 4C-E***) and mimicking its acetylation abolishes TDP-43 nuclear import (***Fig. 3B-F***), three polyclonal antibodies were generated that recognized TDP-43 peptide 77-90 with lysine 82 (K82) acetylation, but not the corresponding unacetylated peptide (***Fig. 5A***). Importantly and as expected (***Fig. 3***), use of each of these antibodies identified increased ac-TDP-43(K82) immunoreactivity on immunoblots of lysates of human neurons following transient, partial proteasome inhibition with BTZ (***Fig. S5***). Moreover, ac-TDP-43(K82) was increased in immunoblotting of lysates of motor cortex in all (six of six) sALS patients tested (albeit with varying levels), while no signal was detected in similar analyses of motor cortex from four non-neurologic disease controls (***Fig. 5B***).

**Figure 5:**
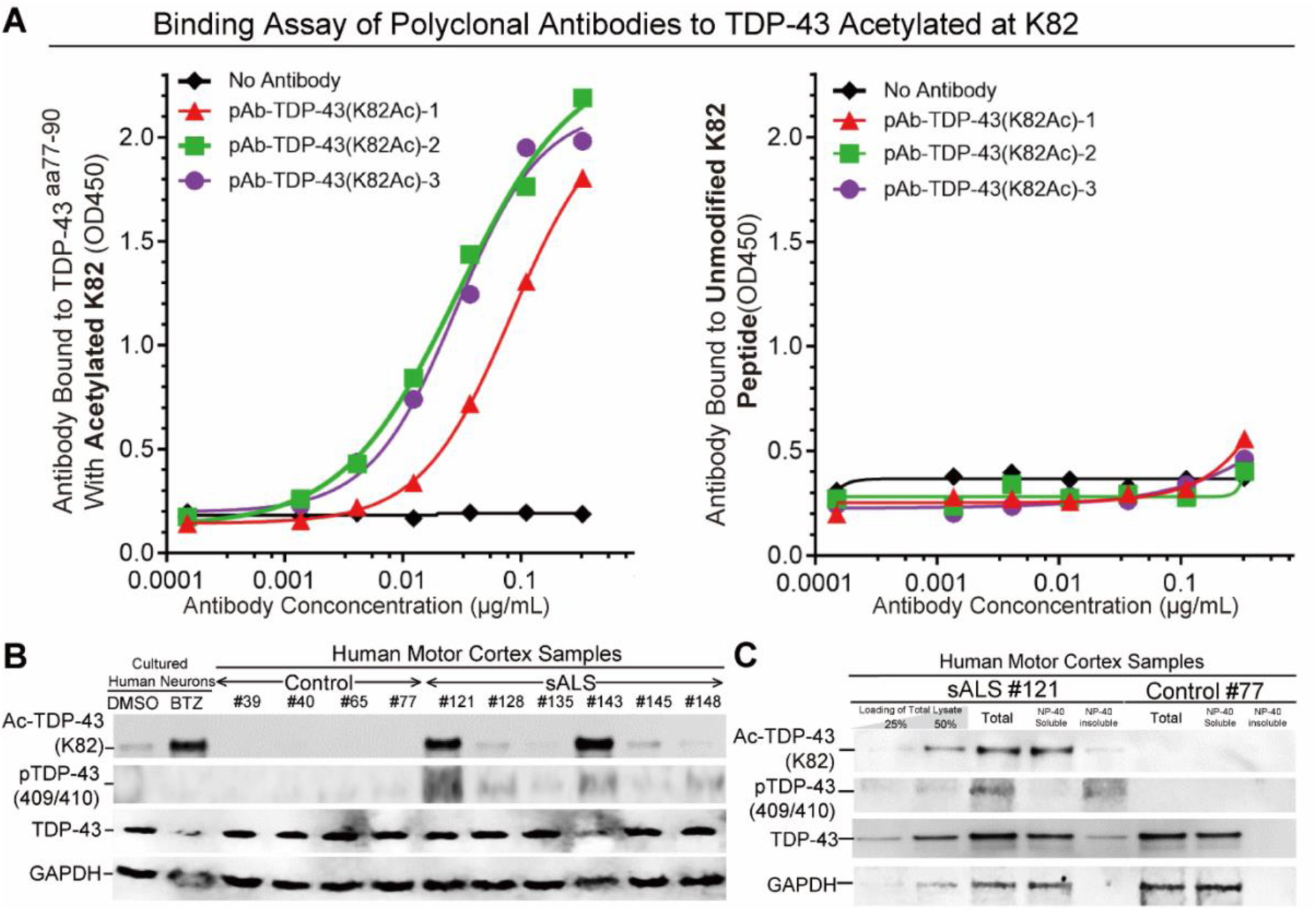
Acetylation of TDP-43 on lysine 82 is increased in sALS postmortem motor cortex. **(A)** Binding specificity of three polyclonal antibodies raised against TDP-43^aa77–90^ peptide acetylated at K82. For each polyclonal antibody, specificity for binding acetylated K82 of TDP-43 was determined by binding to (*left*) TDP-43^aa77–90^ peptide acetylated at K82 or (*right*) the unmodified peptide. **(B)** Levels of TDP-43 acetylated at K82 and phosphorylated at TDP-43 S409/410 determined by immunoblot using antibodies identifying TDP-43, TDP-43 with K82 acetylation, and TDP-43 phosphorylation on S409/S410 in (*left two lanes*) lysates of human iPSC-derived human neurons treated with BTZ or not and (*lanes 3-10*) human fresh frozen motor cortex from (*lanes 3-6*) four non-neurological controls or (*lanes 7-12*) six sALS motor cortices. GAPDH is a loading control. **(C)** Immunoblot using antibodies against TDP-43^aa77–90^ peptide acetylated at K82, TDP-43 with S409/S410 phosphorylation, and total TDP-43 levels in total cortical lysates, NP-40 soluble, and NP-40 insoluble fractions of a sALS individual (#121) and a non-neurological individual (#77). GAPDH serves as a loading control for total cell lysates and NP-40 soluble samples.

sALS samples with the highest ac-TDP-43(K82) signal also had higher phosphorylated TDP-43 (***Fig. 5B***). As expected^31^, phosphorylation of TDP-43 was only detected in the non-ionic detergent insoluble fraction of human freshly frozen motor cortex samples of sALS patients, but not in control subjects. In contrast, immunoreactivity for ac-TDP-43(K82) was identified in both soluble and insoluble fractions of sALS patient cortices, suggesting that acetylation at lysine 82 may be an earlier event than phosphorylation in the pathological cascade driving TDP-43 proteinopathy (***Fig. 5C***).

## Discussion

Our findings demonstrate that TDP-43 nuclear localization is abrogated by lysine 82 acetylation within the TDP-43 cNLS, a process regulated by the opposing activities of acetyltransferases and deacetylases which can be targeted therapeutically^32^. The TDP-43 NLS belongs to the bipartite NLS class with two binding motifs (residues 81-87 and 94-100), with each motif binding to a pocket within importin-α^19^. Our mutagenesis studies confirmed the presence of functional bipartite NLS sequences in TDP-43^15^, since alteration of basic residues in either or both motif (K82A/R83A/K84A and/or K95A/K97A/R98A) led to TDP-43 accumulation in the cytoplasm.

Importantly, using a comprehensive set of single, double or multiple lysine-to-arginine mutants, we determined that lysine K82 is required for stable binding to importin-α1, with optimal binding also relying on a lysine (either K95 or K97) in the other binding pocket. Our findings align with previous reports that had suggested that the TDP-43 NLS belongs to the bipartite NLS class^15,19^, but challenges an earlier hypothesis^33^ that lysine within the NLS regulates importin-α-cargo interactions primarily through its positive charge. Our results show that even a subtle mutation—replacing the positively charged lysine with a similarly charged arginine— was sufficient to disrupt the interaction between TDP-43 and importin-α1. This suggests that any post-translational modification on K82 could lead to TDP-43 mislocalization to the cytoplasm.

Furthermore, using newly made antibodies that specifically recognize lysine acetylation at K82 in TDP-43, we have identified that K82 acetylation is increased in all sALS postmortem motor cortices examined. Our observation is supported by a mass spectrometry analysis of TDP-43 post translational modifications which found K82 acetylation in one out of two ALS postmortem samples^34^. The increase in K82 acetylation was not uniform across the sALS samples we analyzed but correlated with the TDP-43 proteinopathy load, as indicated by the concurrent increase in phosphorylated TDP-43. Additional neuropathological studies are needed to determine whether acetylation at K82 also increases in other TDP-43 proteinopathies including FTD^1^, LATE^7^, and AD^2–6^. The variability in K82 acetylation among sALS patients suggests potential avenues for developing biomarkers that could guide therapeutic strategies targeting TDP-43 acetylation.

Our data support a model in which acetylation initiates and drives TDP-43 proteinopathy. Acetylation at lysine 82 (and K79 and 84 to lesser extents) can initiate TDP-43 nuclear depletion and loss of function through decreased interaction of TDP-43 with its importin receptor. It is possible that differential lysine acetylation events occur in cytosolic and nuclear TDP-43 pools. Indeed, antibodies recognizing acetylated lysine 145 in TDP-43 stained neuropathological inclusions^9^. Hence, it is likely that following K82 acetylation, the cytoplasmically accumulated TDP-43 is further acetylated at lysines K136^35^, K145^9^ and/or K192^9^ which have been reported to decrease RNA binding capacity, increase phase separation^35,36^, aggregation^9,35^, and/or hyperphosphorylation at S409/410 outside the NLS^9,35^, each of which is a key feature of TDP-43 proteinopathies.

## Materials and Methods

### Plasmids

The information for all plasmids used in this paper is listed in Supplementary Table S1. In brief, the entry vector (pST001) was used as a backbone for all the lentiviral vectors. Shortly, traditional cloning methods using double-restriction digestion followed by ligation or Gibson assembly were used to produce the different plasmids. All plasmids in this manuscript will be deposited to Addgene at the time of publication.

### Human neuronal culture derived from induced pluripotent stem cell (iPSC)

The human iPSCs used in this study were a kind gift of Michael Ward, which were previously engineered to express mouse neurogenin-2 (NGN2) under a doxycycline-inducible promoter integrated at the AAVS1 safe harbor in the WTC11 background^37^. iPSCs cells were plated onto Matrigel (Corning, cat. no. 354277) coated plates in Essential 8 Medium (E8; Thermofisher Scientific, cat. no. A1517001) and 10 μM ROCK inhibitor (RI; Y-27632; Selleckchem, cat. no. S1049). iPSCs were maintained in an incubator at 37 °C with 5% CO2 and fed every 1–2 days as needed. Cells were split using EDTA (0.5 mM; Life Technologies, cat. no. 15575020) for routine passaging. Medium was supplemented with 10 μM RI to promote survival during passaging. For neuronal differentiation, iPSCs were plated on day 0 onto a 15-cm plate in N2 medium (knockout Dulbecco’s modified Eagle’s medium (DMEM)/F12 medium; Life Technologies Corporation, cat. no. 12660012) with N2 supplement (Life Technologies, cat. no. 17502048), 1× GlutaMAX (Thermofisher Scientific, cat. no. 35050061), 1× MEM nonessential amino acids (NEAA) (Thermofisher Scientific, cat. no. 11140050), 10 μM ROCK inhibitor (Y-27632; Selleckchem, cat. no. S1049), and 2 μg/ml doxycycline (Clontech, cat. no. 631311). N2 medium was changed once a day for two more days. On day 3, cells were replated onto freshly prepared dishes coated with poly-L-ornithine (PLO; 0.1 mg/ml; Sigma, cat. no. P3655-10MG) and laminin (Sigma, cat. no. L2020-1MG) as follows. Cells were washed with PBS, dissociated with accutase for 10 min at 37 °C, washed and plated in i3Neuron culture medium: Neurobasal™ Medium (Gibco, cat. no. 21103049) supplemented with 1× B27 Plus Supplement (ThermoFisher Scientific, cat. no. A3582801), 10 ng/ml BDNF (PeproTech, cat. no. 450-02), 1 mg/ml mouse laminin (Sigma, cat. no. L2020-1MG), and 2 μg/ml doxycycline (Clontech, cat. no. 631311). Neurons were fed on day 6 during a half-medium change and collected on day 7. For neurons cultured beyond 7 days, half-medium changes were conducted two times per week.

### Cell line culture, transfection and infection

The cell lines used in this paper are HEK293T (ATCC: CRL-11268), and SH-SY5Y (ATCC: CRL-2266). Routine maintenance of these model cell lines follows the guideline posted on ATCC. In brief, HEK293T cells were cultured in complete DMEM supplemented with 10% Fetal bovine serum. SH-SY5Y cells were cultured in DMEM/F12 supplemented with 10% Fetal bovine serum.

Transient transfection was performed for HEK293T cells and at 60% confluency, by using transfection reagent TransIT X2 (Mirus, Cat. no. MIR6000) and following standard protocol posted on the product page of TransIT X2. In brief, serum free DMEM was used to dilute DNA, and then TransIT X2 was added to the mix. After 15 minutes, the final mix was added dropwise to attached cells. The culture medium was changed after 12∼24 hr post transfection. For lentiviral infection, lentiviruses were packed using the 2nd generation packaging system. Briefly, 0.5 x106 per well of 293T cells were seeded in a 6-well plate. For lentiviral transfection, 2.5 µg of the lentiviral plasmid (constructs are labelled as lentivirus in Key Resources Table), 1.25 µg of pMD2.G and 0.625 µg of psPAX2 were inoculated to each well using Mirus transIT-X2 transfection reagent (Mirus). Culture medium was changed to fresh medium at 4∼24 hr post transfection. Two days after transfection, the culture medium was filtered through a 0.45 µm syringe filter to generate the viral supernatant. The viral supernatant containing 10∼50 µg/mL protamine sulfate was added to SH-SY5Y cells for infection. The viral supernatant was removed 24 hr after infection, and cells were passaged at least once before selection. Infected cells are selected based on the selection marker encoded by the lentivirus. For SH-SY5Y cells, the concentrations were 400 µg/mL for neomycin, 10 µg/mL for blasticidin-S and 3 µg/mL for puromycin. Detailed guides and protocols posted can be found on the Addgene website: https://www.addgene.org/protocols/lentivirus production / https://www.addgene.org/guides/lentivirus/

### Central Nervous System (CNS) tissues

Human tissues were obtained from the UCSD ALS tissue repository that was created following HIPAA-compliant informed consent procedures approved by Institutional Review Boards (either Benaroya Research Institute, Seattle, WA IRB# 10058 or University of California San Diego, San Diego, CA IRB# 120056). Spinal cord and brain tissues were acquired using a short-postmortem interval acquisition protocol usually under 6 h. Tissues were immediately dissected in the autopsy suite, placed in labeled cassettes, and fixed in neutral buffered formalin for at least 2 weeks before being dissected and paraffin-embedded for indefinite storage. For this study, we evaluated from 4 control cases, 6 sALS cases (Supplementary Table S2).

### Chemicals and cell treatments

Doxycycline (DOX Sigma-Aldrich, Cat. no. D9891) was used to induce gene expression via the TetON3G promoter. Bortezomib (BTZ, ApexBio, Cat. no. A2614), MG132 (Selleckchem, Cat. no. S2619) and Marizomib (Selleckchem, Cat. no. S7504) were used to treat to SH-SY5Y cells and iPSC-derived cortical neurons to decrease proteasome activity.

### Cell Survival Assay

Cell survival assay was performed using the CellTiter-Glo Luminescent Cell Viability Assay kit. A CellTiter-Glo Assay (Promega) was performed according to the manufacturer’s instructions. Luminescence was recorded with a Tecan GENios Pro plate reader.

### Proteasome Activity Assay

Proteasome activity assay was performed using the Proteasome-Glo™ reagent (Promega, cat. no. G8621) and coupled with a CellTiter-Glo Assay (Promega, cat. no. G9241) to control for cell viability. Both assays were performed according to the manufacturer’s instructions. Luminescence was recorded with a Tecan GENios Pro plate reader.

### Confocal Microscopy

For immunofluorescence, cells were first fixed with 4% PFA for 10 minutes at room temperature (RT), washed twice with PBS, and permeabilized by using 0.5% Triton X-100 in PBS. Then cells were incubated with blocking buffer (2% BSA, 0.1% Triton X-100 in PBS) for 30 minutes at RT. Primary antibodies were diluted in the blocking buffer, and cells were incubated with primary antibody over night at 4°C. Antibodies and dilutions are provided in Supplementary Table S3. Then cells were washed 3 times at RT with PBS. Secondary antibodies were dilute in PBS at 1:500 dilution and incubated for 45 minutes at RT, covered from light. After the incubation with secondary antibodies, cells were washed 3 times at RT with PBS and stained with DAPI (Tocris, Cat. no. 5748) at 1μg/mL in PBS. Then cells were mounted with ProLong Gold (Thermofisher, Cat. no. P10144). The slides were cured at RT overnight in dark. Spinning disk confocal Yokogawa X1 confocal scanhead mounted to a Nikon Ti2 microscope with a Plan apo lambda 100 × oil NA 1.45 objective and Plan apo lambda 60 × oil na 1.4 objective.

### Live-Cell Imaging

For live-cell imaging, either SH-SY5Y cells or human iPSC-derived cortical neurons expressing the fluorescently tagged protein of interest was plated on 96-well, glass-like, polymer-bottom plates (Cellvis, cat. no. P96-1.5H-N) and treated as indicated. Live-cell images were acquired for 0–48 h on the CQ1 benchtop spinning-disk confocal high-content analysis system (Yokogawa) (CQ1 software v.1.05.01.02). Cells were imaged using a ×40 or ×60 dry objective and maintained under humidified conditions at 37 °C in the presence of 5% CO_2_ during imaging.

### Nucleus/Cytoplasmic fractionation

Nucleus/cytoplasmic fractionation was performed following the protocol of Abcam Nuclear Extraction Kit (abcam, Cat. no. ab113474). In brief, 70-80% confluency cultured cells on a 10 cm dish (approximately 2-5 × 10^6 cells), were washed twice with PBS (phosphate-buffered saline), scraped, transferred to a 15 mL conical tube, and centrifuged at 1000 rpm for 5 minutes. After centrifugation, the supernatant was discarded and the cell pellet was resuspended in 100 µL of 1 x Pre-Extraction Buffer, incubated on ice for 10 minutes, vigorously vortexed for 10 seconds, and centrifuged at 12,000 rpm for 1 minute. The supernatant was collected as the cytoplasmic extract and the pellet as nuclear fraction. The cytoplasmic and nuclear protein fraction were quantified and used for subsequent applications.

### Quantitative mass spectrometry for the nuclear proteome

#### Nuclear isolation

Nuclear and cytoplasmic fractions were isolated from human iPSC-derived cortical neurons using the NE-PER Nuclear and Cytoplasmic Extraction Reagents (Thermo Scientific cat. no. 78833). Cells were harvested via centrifugation at 500 × g for 5 minutes and washed with PBS. The cell pellet was resuspended in ice-cold Cytoplasmic Extraction Reagent I (CER I) and incubated on ice for 10 minutes. Cytoplasmic Extraction Reagent II (CER II) was then added, followed by brief vertexing and centrifugation at 16,000 × g for 5 minutes to separate the cytoplasmic extract. The nuclear pellet was subsequently resuspended in Nuclear Extraction Reagent (NER) and subjected to intermittent vertexing on ice for 40 minutes. The nuclear extract was collected following centrifugation at 16,000 × g for 10 minutes and pellet was discarded. Nuclear extracts were stored at -80°C until analysis. Protease inhibitors were included in the CER I and NER to maintain protein integrity.

#### Protein digestion and TMT labelling

After three washes with wash buffer 1 (100 mM TEAB pH 8.0, 4 M urea and 0.5% SDS) and four washes with wash buffer 2 (100 mM TEAB pH 8.0 and 4 M urea), nuclear fractions were resuspended in 100 mM TEAB and 2 M urea supplemented with 10 ng µl−1 trypsin and 5 ng µl−1 lys-C for 1-hr pre-digestion at 37 °C on a thermomixer with shaking at 1,000 rpm. The pre-digested products were collected and an additional 14-hr digestion with 10 ng µl−1 trypsin at 37 °C. The digested peptides from each sample were labelled with TMT six-plex labelling reagents (Thermo Scientific, 90061) following the manufacturer’s instructions. Briefly, the TMT reagents were solubilized in anhydrous acetonitrile and added to peptides from each sample according to the labelling. Following incubation at room temperature (RT) for 1 h, 5% hydroxylamine was added to the samples and incubated for a further 15 min to quench the reaction. Equal volumes of peptides from each sample in the same group were pooled and dried in a SpeedVac to remove the acetonitrile. The samples were acidified with formic acid (final concentration of 1%) and desalted using Pierce C18 spin columns (89870).

#### Liquid chromatography with mass spectrometry analysis

The TMT-labelled samples were analyzed on a Orbitrap Eclipse mass spectrometer (Thermo Scientific). The samples were injected directly onto a 25 cm, 100 μm inner diameter column packed with BEH 1.7 μm C18 resin (Waters) and separated at a flow rate of 300 nl min−1 on an nLC 1200 system (Thermo Scientific). Buffers A and B were 0.1% formic acid in 5% acetonitrile and 80% acetonitrile, respectively. A gradient of 0–25% Buffer B over 75 min, an increase to 40% Buffer B over 30 min, an increase to 100% Buffer B over another 10 min and holding at 100% Buffer B for 5 min was used for a total run time of 120 min.

Peptides were eluted directly from the tip of the column and nano-sprayed directly into the mass spectrometer by application of 2.5 kV voltage at the back of the column. The Eclipse was operated in data-dependent mode. Full MS1 scans were collected in the Orbitrap at 120 k resolution. The cycle time was set to 3 s and within these 3 s the most abundant ions per scan were selected for collision-induced dissociation tandem mass spectrometry in the ion trap. MS3 analysis with multi-notch isolation (SPS3) was utilized for detection of TMT reporter ions at 7.5 k resolution^38^. Monoisotopic precursor selection was enabled, and dynamic exclusion was used with an exclusion duration of 60 s.

#### Analysis of quantitative mass spectrometry data

Raw data were processed using Rawconverter ^39^ to extract MS2 and MS3 spectra with a correction of each precursor ion peak to its monoisotopic peak when appropriate. The MS2 and MS3 mass spectrometry spectra were searched against a complete human protein database downloaded from UniProt. The search parameters used were: precursor mass tolerance of 50 ppm, fragment ion tolerance of 500 ppm for collision-induced dissociation spectra and of 20 ppm for higher-energy C-trap dissociation spectra, minimum peptide length of six amino acids, static modifications for carbamidomethylation of cysteine and TMT tags on lysine residues and peptide N termini (+229.162932 Da). The identified peptide-spectrum matches (PSMs) were filtered to a false-detection rate of ≤1% at a PSM level with DTASelect2.

The false-detection rate was calculated based on the number of PSMs that matched to sequences in the reverse decoy database. TMT quantification of reporter ions from MS3 spectra was performed using Census2 with a filter of over 0.6 for isobaric purity. The normalized intensity based on weighted normalization was used to calculate the ratio of reporter ions corresponding to the indicated groups. The ratios of each protein from three forward labelling groups and three reverse labelling groups (Supplementary Table 1) were used to calculate P values using a one-sample Student’s t-test. The volcano plot was generated using the R package.

### Immunoblotting

Tissue and cell samples were lysed in RIPA buffer (containing 50 mM Tris-HCl pH 7.4, 150 mM NaCl, 0.1% SDS, 1% NP-40, 0.25% sodium deoxycholate, 1 mM sodium fluoride, and 1 mM Na3VO4), supplemented with an EDTA-free protease inhibitor cocktail (Roche Life Science, Cat. no. 11873580001) and a phosphatase inhibitor cocktail (Roche Life Science, Cat. no. 04906837001). Protein concentrations were quantified using the Bradford method. Protein samples were separated by SDS-PAGE, followed by transfer onto polyvinylidene difluoride membranes. The membranes were blocked using 5% skim milk (BD Difco, Cat. no. 232100) in PBS containing 0.05% Tween-20 (0.05% TBST). Subsequently, they were incubated with primary antibodies at 4°C for 24 hr (Supplementary Table S4). After this incubation, the membranes were probed with horseradish peroxidase (HRP)-conjugated secondary antibodies for 1 hr, and protein bands were visualized using an enhanced chemiluminescence (ECL) detection Kit (Vazyme, Cat. no. E423-02).

### Immunoprecipitation (IP) and Co-Immunoprecipitation (Co-IP)

For post-translational modification (PTM) MS analysis, iPSC-derived cortical neurons or SH-SY5Y cells that endogenously express TDP-43 or stably express TDP-43 variants (clover-tagged) were collected and stored overnight as cell pellets in liquid nitrogen. The cell pellets were lysed using RIPA buffer supplemented with protease inhibitors (Roche Life Science, Cat. no. 11873580001), PhosSTOP (Roche Life Science, Cat. no. 04906837001), and benzonase (Millipore, cat. no. 70664-3). To shear the genomic DNA, the cell lysate was sonicated twice for 2 seconds at 35% amplitude. After a 30-minute incubation on ice, magnetic beads specific to different antigens (Supplementary Table S5) were washed twice with complete RIPA buffer. For the IP experiment, 90% of the supernatant was added to 50 μl of magnetic beads and incubated at 4°C for 12 hr. The supernatant was then removed, and the beads were washed at least ten times with complete RIPA buffer at 4°C. The beads were then prepared for MS spectrometry analysis.

For the Co-IP experiment, cell pellet preparation follows the same protocol as for IP. The main difference is that Co-IP uses IP lysis buffer (Thermo Fisher, cat. No. 87787) without SDS or other strong detergents, along with PhosSTOP, protease inhibitors, and benzonase. 90% of the supernatant was added to 50 μl of affinity beads and incubated at 4°C for 1-3 hr. Proteins bound to the beads were released by adding 6x Laemmli SDS sample buffer and boiling for 10 minutes, preparing them for subsequent immunoblotting.

### Silver Staining

ProteoSilver™ Silver Stain Kit (Sigma, cat. no. PROTSIL1) was used for silver staining following the kit’s standard protocol. Briefly, after SDS-PAGE, the gel was fixed in a solution of 50% ethanol, 10% acetic acid, and 40% water, and then washed with water. Sensitization with thiosulfate was performed to enhance contrast before the gel was incubated with silver nitrate to bind proteins. After washing away the excess of silver, the gel was developed with formaldehyde and sodium carbonate to visualize proteins as dark bands, and the development was stopped with acetic acid followed by a final water rinse prior imaging.

### PTM Detection by Mass Spectrometry

After isolating TDP-43 using the IP protocol, beads containing TDP-43 were sent to the UCSD mass spectrometry core. TDP-43 was digested using trypsin and chymotrypsin independently, and then peptides were combined and enriched using titanium dioxide chromatography. The enriched peptides were then analyzed by liquid chromatography-tandem mass spectrometry (LC-MS/MS), using either data-dependent or data-independent acquisition methods to identify and quantify the modifications. Data analysis was conducted with software tools (MaxQuant or Proteome Discoverer) to identify the peptides and their PTMs.

### TDP-43 Peptide-Importin-α1-ΔIBB binding assay

The TDP-43-77-110 peptide-Importin-α1-ΔIBB binding assay begins by coating the plates with antigens at a concentration of 8 μg/mL in 1× PBS, with 30 µL per well, and incubating them overnight at 4°C. Following incubation, the plates are washed three times with PBST and then blocked with 5% BSA for 2 hr at room temperature (RT). After another three washes with PBST, Importin-α1-ΔIBB-His (Synthesised by Sanyou. Inc.) in 1% BSA is added to the plates, with 30 µL per well, and incubated for 60 minutes at RT. The plates are then washed three more times with PBST before adding Anti-6*His-HRP (Proteintech, cat. no. HRP-66005) at a 1:4000 dilution in 1% BSA and incubating for 60 minutes at RT. Finally, the plates are washed six times with PBST, TMB (3,3’,5,5’-tetramethylbenzidine) is added to develop the reaction, and the reaction is stopped with stop solution before reading the OD450.

### Generation of TDP-43 lysine 82 acetylation antibody

Polyclonal antibodies to TDP-43 acetylated at K82 were generated by Sanyou. Inc. A peptide corresponding to TDP-43 residues 77-90 with or without acetylation at K82, was synthesized and conjugated to the carrier protein KLH to enhance immunogenicity. Rabbits were immunized with this peptide-carrier conjugate, with an initial injection followed by regular booster injections to stimulate the immune response. Antibody titers were monitored via ELISA. Once sufficient antibody levels were reached, blood was collected from the rabbits, allowed to clot at room temperature, and the serum was separated by centrifugation. Following affinity enrichment, antibodies specifically recognizing TDP-43 acetylation at lysine 82 were isolated and validated through ELISA.

### ELISA-based binding assays to validate antibody specificity

ELISA assay begins by coating the plates with TDP-43 residues 77-90 containing or not containing K82 acetylation at a concentration of 8 μg/mL in 1× PBS, with 30 µL per well, and incubating them overnight at 4°C. Following incubation, the plates are washed three times with PBST and then blocked with 5% BSA for 2 hr at room temperature (RT). After another three washes with PBST, three polyclonal antibodies generated that recognized TDP-43 peptide 77-90 with lysine 82 (K82) acetylation (Synthesised by Sanyou. Inc.) underwent series dilution in PBST. 1% BSA was added to the plates, with 30 µL per well, and incubated for 60 minutes at RT. The plates are then washed three more times with PBST before adding Anti-rabbit-HRP (Proteintech, cat. no. SA00001-2) at a 1:4000 dilution in 1% BSA and incubating for 60 minutes at RT. Finally, the plates are washed six times with PBST, TMB (3,3’,5,5’-tetramethylbenzidine) is added to develop the reaction, and the reaction is stopped with stop solution before reading the OD450.

### TDP-43 Peptide-Importin-α1-ΔIBB binding assay

The TDP-43-77-110 peptide-Importin-α1-ΔIBB binding assay begins by coating the plates with antigens at a concentration of 8 μg/mL in 1× PBS, with 30 µL per well, and incubating them overnight at 4°C. Following incubation, the plates are washed three times with PBST and then blocked with 5% PBSM for 2 hr at room temperature (RT). After another three washes with PBST, Importin-α1-ΔIBB-His (Synthesised by Sanyou. Inc.) in 1% PBSM is added to the plates, with 30 µL per well, and incubated for 60 minutes at RT. The plates are then washed three more times with PBST before adding Anti-6*His-HRP (Proteintech, cat. no. HRP-66005) at a 1:4000 dilution in 1% PBSM and incubating for 60 minutes at RT. Finally, the plates are washed six times with PBST, TMB (3,3’,5,5’-tetramethylbenzidine) is added to develop the reaction, and the reaction is stopped with stop solution before reading the OD450.

### RNA extraction, qRT-PCR quantification, and RT-PCR

Cultured IPSC-derived cortical neurons were directly lysed in 1ml Trizol reagent (Thermo, cat. no. 15596026), followed by chloroform addition (200 μl) and RNA extraction accordingly. Total RNA was quantified on a Nanodrop spectrophotometer and 1 μg was taken forward to first-strand cDNA synthesis with SuperScript-III reverse transcriptase kit (Thermo, cat. no. 18080044) and oligo dT priming per manufacturer instructions. For qRT-PCR, cDNA was diluted to 1 ng/μl, and 4 ng were loaded into a 10 μl reaction (BioRad). Three technical replicates were assayed per biological sample and analyzed on a C1000 thermocycler with a 384-well qPCR reaction module (Bio-Rad) using primers/probes detailed in Supplementary Table S5. Human *GAPDH* gene was used as endogenous control. Relative expression for each gene was calculated from delta-delta-Cq data.

### Statistical tests

Statistical analyses were performed using Prism 8 software by GraphPad. Each data point represents an independent biological replicate (distinct wells of independently treated cells or individual tissue donors). For comparisons between two groups, statistical significance was assessed using two-tailed Student’s t-tests. For comparisons involving three or more groups, one-way ANOVA with Tukey’s correction or Chi-squared tests with Yates’ correction were applied as noted in the figure legends. Significance levels were denoted as follows: ****P < 0.0001; ***P < 0.001; **P < 0.01; *P < 0.05; ns, P > 0.05. Error bars represent SEM unless stated otherwise.

---

**Supplementary Figure 1.**
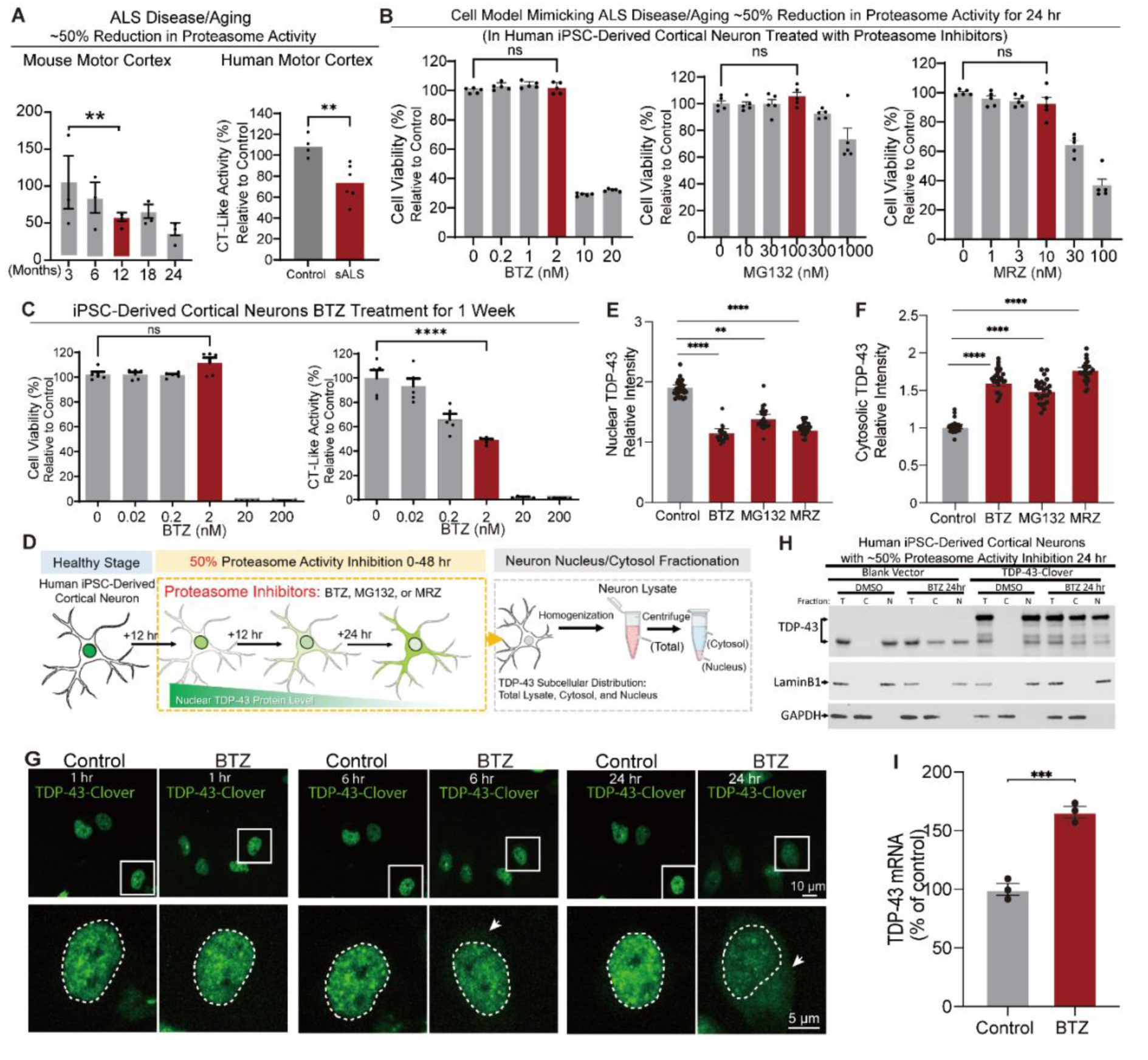
Partial proteasome inhibition results in TDP-43 mislocalization and loss of function. **(A)** Evaluation of CT-like proteasome activity in cell lysates from human or mouse fresh frozen motor cortex samples. Human samples are from n=4 non-neurological control individuals (Control) and n=6 sporadic ALS patients (sALS). Mouse samples were collected from 3, 6, 12, 18, 24 month old mice. **(B)** Cell viability analysis determined by ATP levels in cell lysates from human induced pluripotent stem cell (iPSC)-derived cortical neurons exposed to varying doses of proteasome inhibitor BTZ for 24 hr. **(C)** CT-like proteasome activity and cell viability analysis for human induced pluripotent stem cell (iPSC)-derived cortical neurons exposed to varying doses of proteasome inhibitor BTZ for 1 week. **(D)** Schematic of experimental design to identify TDP-43 subcellular localization after proteasome inhibitor mediated decrease in proteasome activity in human iPSC-derived cortical neurons. **(E)** Quantification of the intensity of nuclear TDP-43 in Fig. 1C relative to control. **(F)** Quantification of the intensity of cytosolic TDP-43 relative to control in Fig. 1C. **(G)** Representative live-cell images of TDP-43-Clover in human iPSC-derived cortical neurons after partial proteasome inhibition mediated by addition of BTZ (2 nM) for 0-24 hr. **(H)** Immunoblot to determine TDP-43 and TDP-43-Clover protein levels in total cell lysate (T), nucleus (N), and cytosol (C) from human iPSC-derived cortical neurons. Neurons were transfected to express TDP-43-Clover or an empty plasmic vector for 24 hr, followed by partial, BTZ (2 nM) mediated proteasome inhibition for 24 hr. Immunoblots for lamin B1 and GAPDH are used as loading controls for nuclear and cytosolic fractions. **(I)** TDP-43 mRNA levels determined by RT-PCR in lysates of human iPSC-derived cortical neurons with or without complete proteasome inhibition following BTZ addition (20 nM) for 24 hr.

**Supplementary Figure 2.**
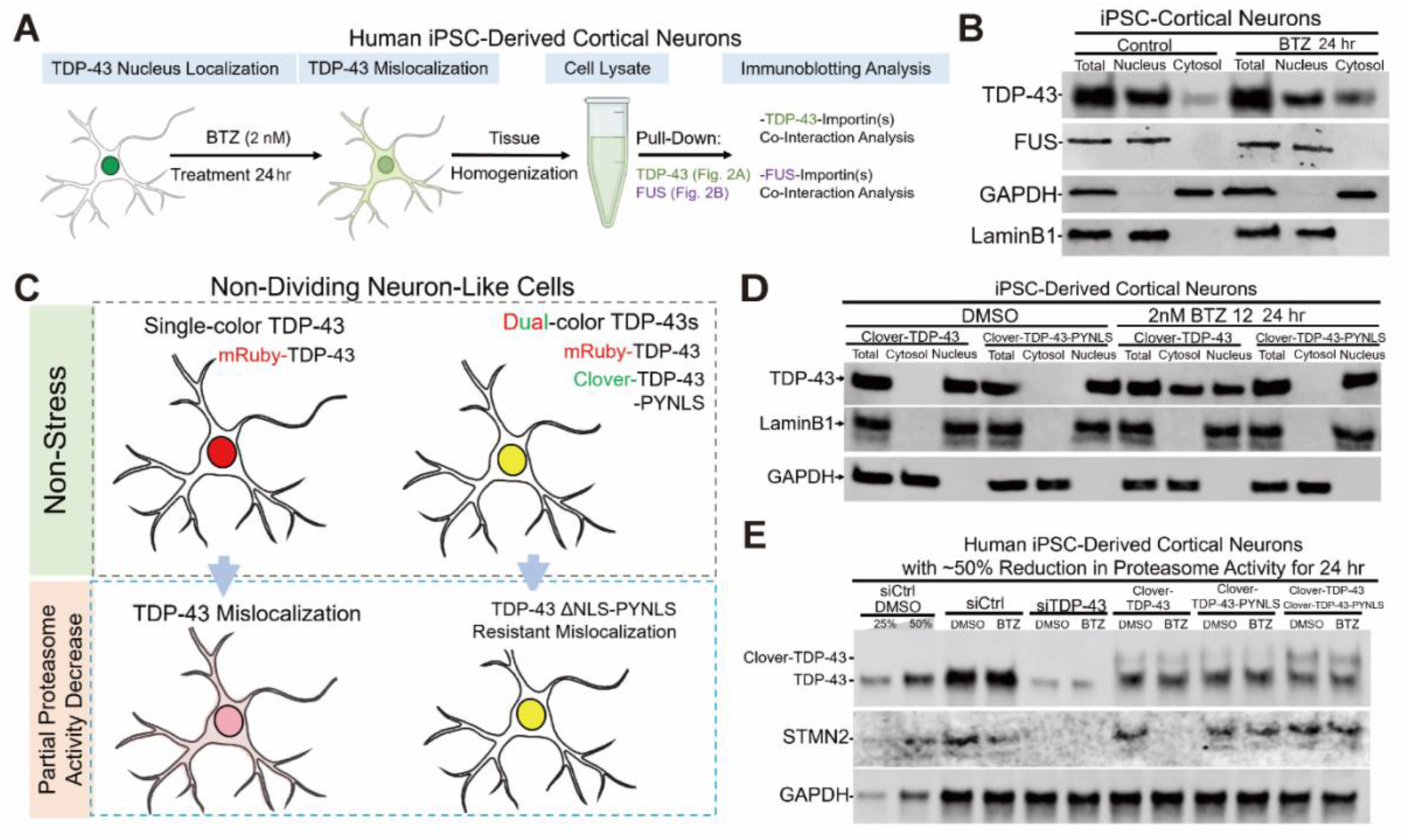
TDP-43 mislocalization and loss of function after partial proteasome inhibition is rescued by replacing the TDP-43 cNLS with the PY-NLS from FUS. **(A)** Schematic of analysis of importin-α1/TDP-43 or importin-β2/FUS interactions determined by immunoprecipitating TDP-43 or FUS from iPSC-derived cortical neurons after BTZ (2 nM) treatment for 24 hr. **(B)** Immunoblot to determine TDP-43 protein levels in total cell lysates, nuclear, or cytosol fractions of human iPSC-derived cortical neurons with or without partial BTZ (2 nM) mediated proteasome inhibition for 24 hr. Immunoblots for lamin B1 and GAPDH provide loading controls for nuclear and cytosolic fractions**. (C)** Schematic of cell fractionation of human neuron-like, SH-SY5Y cells expressing TDP-43 or TDP-43-PY-NLS, with and without partial proteasome inhibition for 6 hr. **(D)** Immunoblot to determine protein levels of Clover-TDP-43 and Clover-TDP-43-PYNLS within total cell lysates, nuclear, or cytosol fractions of human iPSC-derived cortical neurons with or without partial proteasome inhibition from BTZ (2 nM) addition for 24 hr. Immunoblots for lamin B1 and GAPDH provide loading controls for nuclear and cytosolic fractions. **(E)** Immunoblot detection stathmin-2 protein level in human iPSC-derived cortical neurons expressing Clover-TDP-43 or Clover-TDP-43-PY-NLS with or without partial BTZ (2 nM) mediated proteasome inhibition for 24 hr.

**Supplementary Figure 3:**
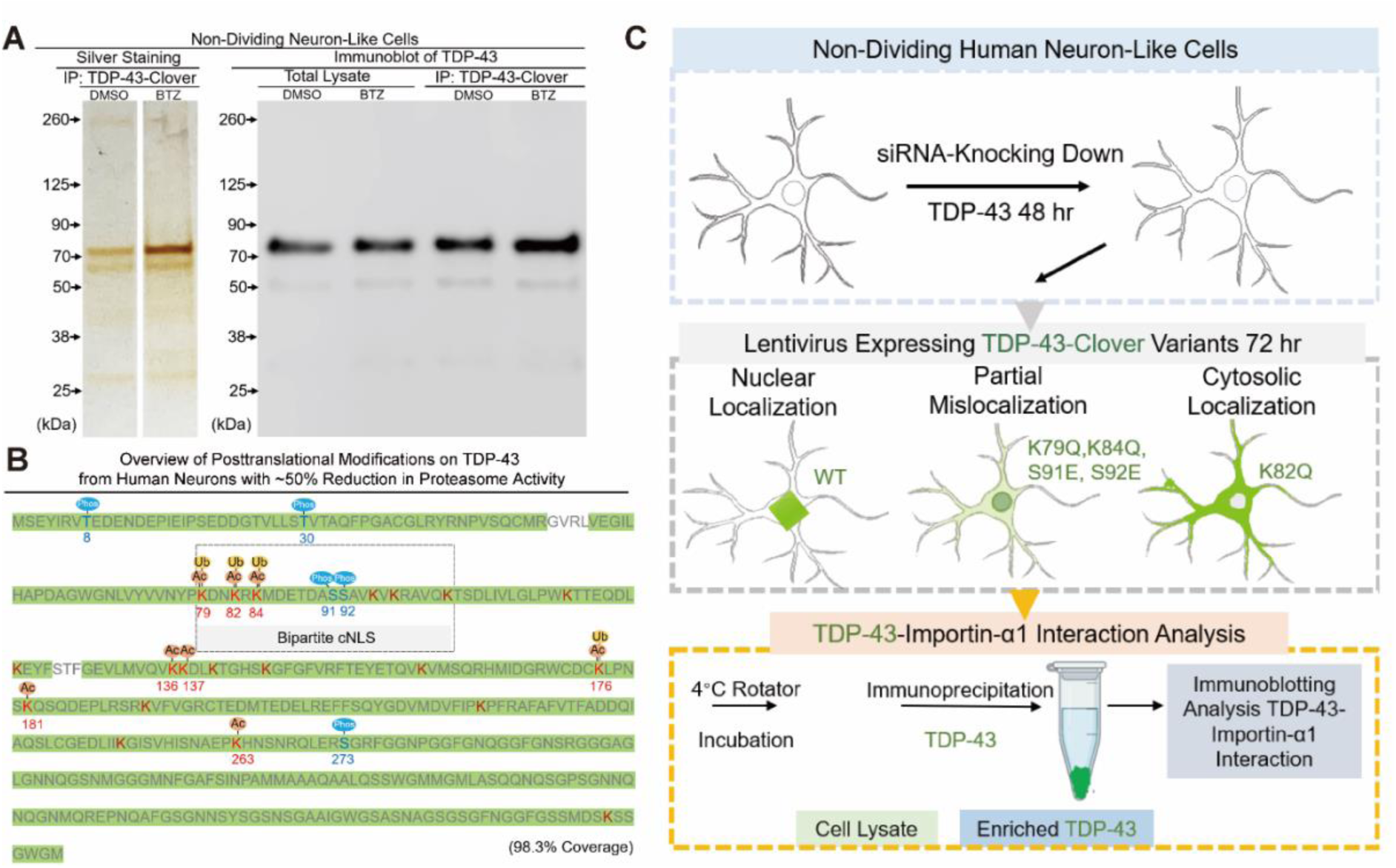
Identification of post-translational acetylation, ubiquitination, and phosphorylation within the TDP-43 cNLS after partial proteasome inhibition. **(A)** (*Left*) Silver stain and (*right*) immunoblot of TDP-43-Clover affinity purified from non-dividing human neuron-like SH-SY5Y cells with homozygous knock-in of TDP-43-Clover at both endogenous TDP-43 alleles. Partial proteasome inhibition was first induced in SH-SY5Y cells by addition (or not) of the BTZ (2 nM) for 24 hr, after which TDP-43-Clover was affinity purified from cell lysates using GFP nanobodies conjugated to magnetic beads, and finally mass spectrometry was used to identify post-translational modifications. **(B)** Summary of the posttranslational modifications of TDP-43 identified in (A). **(C)** Schematic of experimental design to assess functional consequence of mimics of the post-translational modifications identified within the TDP-43 cNLS in (B). SH-SY5Y cells were pre-treated with either control or human TDP-43 siRNA for 48 hr, followed by lentivirus-driven expression of each TDP-43 variant (with C-terminal Clover tag) for 72 hr. Each TDP-43-Clover tagged variant was affinity purified using GFP nanobodies conjugated to magnetic beads and bound importin-α1 was detected by immunoblotting (see Fig. 3C***-E***).

**Supplementary Figure 4:**
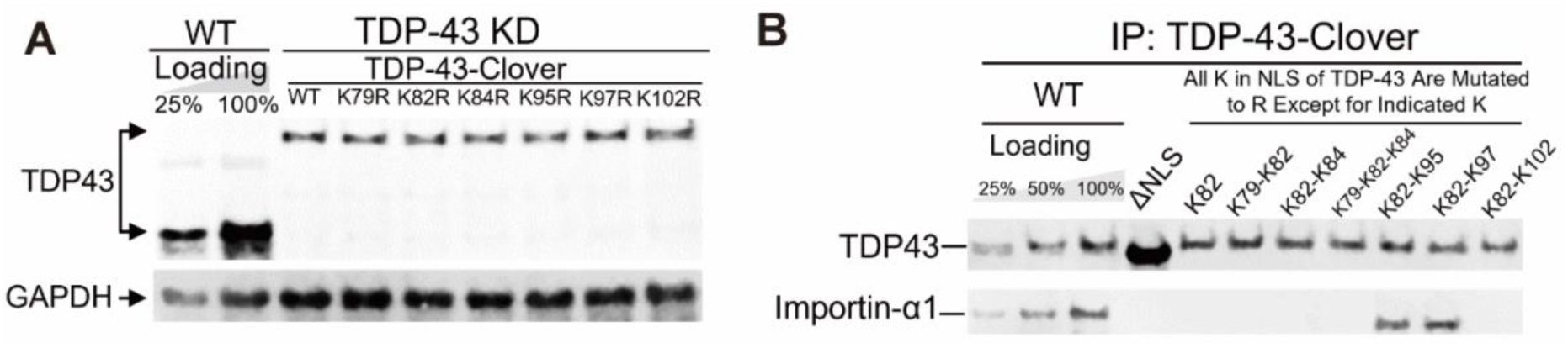
Lysine 82 (K82), together with K95 or K97, within the bipartite cNLS of TDP-43 are necessary and sufficient for TDP-43 binding to importin-α1. **(A)** Immunoblot determination of accumulated level of lysine to arginine (K to R) variants (see ***Fig. 4C-E***) within the TDP-43 cNLS of Clover tagged TDP-43 in non-dividing human neuron-like SH-SY5Y cells treated with either control or human TDP-43 siRNA for 48 hr, followed by lentivirus-mediated expression for 72 hr of each TDP-43 variant carboxy-terminally tagged with Clover. GAPDH was used as a loading control. **(B)** Affinity purification of TDP-43-Clover from lysates of the cells in (A) was performed with GFP nanobody-magnetic beads followed by immunoblotting to detect the levels of TDP-43 and importin-α1.

**Supplementary Figure 5.**
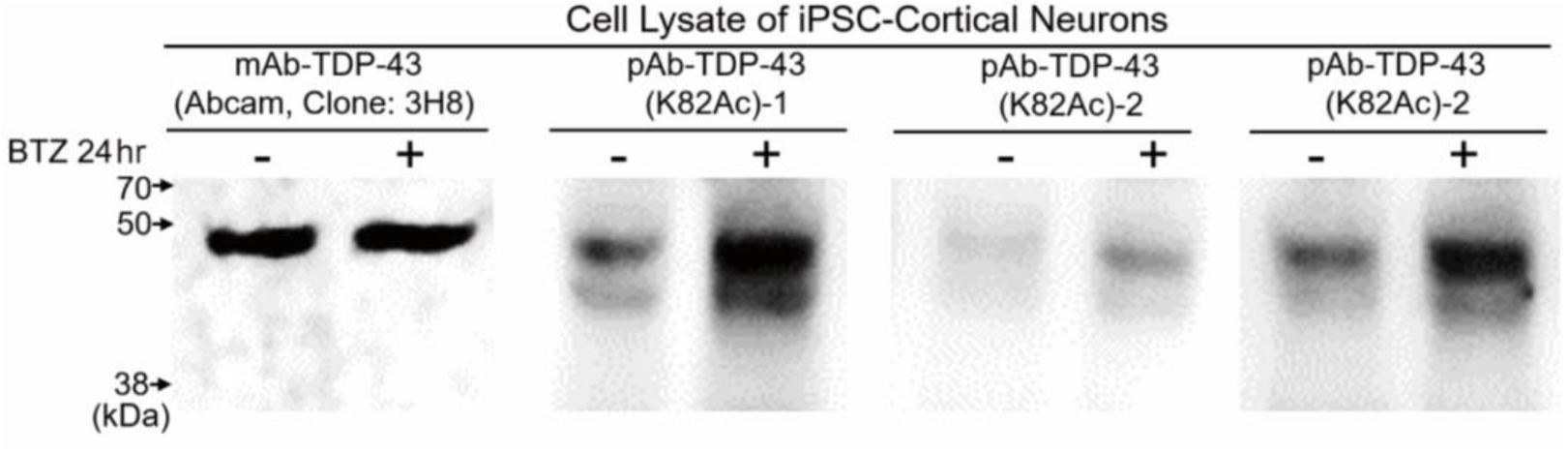
Increased lysine 82 acetylation of TDP-43 in iPSC-derived cortical neurons after partial proteasome inhibition. Immunoblot showing levels of total TDP-43 protein and TDP-43 acetylated at K82 (detected with each of three polyclonal antibodies raised recognizing TDP-43 acetylated at K82) in lysates of human iPSC-derived cortical neurons with or without partial proteasome inhibition produced by addition of BTZ (2 nM) for 24 hr.

